# Diel activity patterns of vector mosquito species in the urban environment: Implications for vector control strategies

**DOI:** 10.1101/2022.08.24.505011

**Authors:** André B. B. Wilke, Adequate Mhlanga, Allisandra G. Kummer, Chalmers Vasquez, Maday Moreno, William D. Petrie, Art Rodriguez, Christopher Vitek, Gabriel L. Hamer, John-Paul Mutebi, Marco Ajelli

## Abstract

Florida and Texas continue to be afflicted by mosquito-borne disease outbreaks such as dengue and West Nile virus disease and were the most affected states by the Zika outbreak of 2016-2017. Mathematical models have been widely used to study the population dynamics of mosquitoes as well as to test and validate the effectiveness of arbovirus outbreak responses and mosquito control strategies. The objective of this study is to assess the diel activity of mosquitoes in Miami-Dade, Florida and Brownsville, Texas, and to evaluate the effectiveness of simulated adulticide treatments on local mosquito populations. To assess variations in the diel activity patterns, mosquitoes were collected hourly for 96 hours once a month from May through November 2019 in Miami-Dade and Brownsville, Texas. We then performed a PERMANOVA followed by the SIMPER method to assess which species contributed the most to the observed differences. Finally, we used a mathematical model to simulate the population dynamics of 5 mosquito vector species to evaluate the effectiveness of the simulated adulticide applications. A total of 14,502 mosquitoes comprising 17 species were collected in Brownsville and 10,948 mosquitoes comprising 19 species were collected in Miami-Dade. *Aedes aegypti* was the most common mosquito species collected every hour in both cities and peaking in abundance in the morning and the evening. Our modeling results indicate that the effectiveness of adulticide applications varied greatly depending on the hour of the treatment. Overall, 9 PM was the best time for adulticide applications targeting all mosquito vector species in Miami-Dade and Brownsville. Our results indicate that the timing of adulticide spraying interventions should be carefully considered by local authorities based on the ecology of mosquito species in the focus area.

## Introduction

The invasion and proliferation of mosquito vector species across new areas directly affect the epidemiology of mosquito-borne diseases, especially in complex socio-ecological urban ecosystems [1–3]. Due to the increase in the presence and abundance of vector mosquito species, major cities in the contiguous United States have been affected by local arbovirus transmission [4,5]. In particular, southern Florida and southern Texas have been historically afflicted by arbovirus outbreaks and were the most affected areas during the Zika virus outbreak of 2016 in the United States [3,6]. Furthermore, in 2020, 6 locally transmitted human cases of dengue and 59 (symptomatic and asymptomatic) infections of West Nile virus disease were reported in Miami-Dade, Florida, and a major dengue outbreak with 65 confirmed locally transmitted cases were reported in the Florida Keys [5,7]. Further epidemiological investigation in Miami-Dade detected the circulation of West Nile virus and dengue serotypes without reported human cases [8]. International travel was severely restricted at that time due to the COVID-19 pandemic indicating local subclinical circulation of dengue and possibly other arboviruses in the United States.

The increase in the number of arboviral infections is most likely due to the increase in the contact rate between mosquito vector species and human hosts [9,10], and even though controlling mosquitoes is considered the most effective way to prevent arbovirus outbreaks, controlling mosquitoes in urban areas is a difficult task [11]. Mosquitoes have very high reproductive potential and can lay hundreds of eggs during their lifetime; under normal conditions, only a selected few survive to adulthood [12–14]. However, due to the lack of or reduced density of natural predators, reduced larval competition and an overabundance of resources in urban areas (including widely available artificial aquatic habitats for larval development and human hosts for blood-feeding) there is a greater number of female mosquitoes laying eggs and a large proportion of immature mosquitoes surviving to adulthood. This leads to large mosquito populations, including vector species, in urban areas [15–17].

Due to this ecological imbalance, mosquito vector populations can reach great numbers in urban areas. One of the most common strategies to control mosquitoes in urban areas is to reduce the number of active host-seeking females by spraying adulticides [18]. Due to current environmental standards, the development of insecticide resistance, and undesired effects on non-target organisms and human populations the application of adulticides should ideally be used under the Integrated Vector Management (IVM) framework [18–22]. However, adulticide spraying is still one of the most important tools for controlling mosquitoes and is widely used worldwide in non-emergency situations, especially in developing countries with limited access to other, more expensive and ecologically friendly, alternatives such as bacterial larvicides and insect growth regulators.

One of the most important elements that separate successful from unsuccessful adulticide spraying is the extent to which the target adult mosquito population is exposed to the adulticide [23]. To effectively reduce a mosquito population adulticide applications must coincide with the period when the mosquitoes are most active (actively flying). On the other hand, adulticide interventions will have only have very limited effect on the resting adult mosquito population especially those in cryptic resting sites. Therefore, knowing the period(s) of the day mosquitoes are most active is of paramount importance for vector control operations and for the prevention and control of local arboviral transmission.

Secondly, to leverage a mathematical model for the dynamics of five arbovirus vector species to evaluate the effectiveness of adulticide applications at different periods of the day and at different frequencies.

## Results

### Descriptive analysis

A total of 14,502 mosquitoes comprising 17 species were collected in Brownsville and 10,948 mosquitoes comprising 19 species were collected in Miami-Dade. *Aedes aegypti* was the most common mosquito species collected every hour of the day in both Brownsville and Miami-Dade. It was also the most abundant species, totaling 7,024 and 4,444 collected specimens in Brownsville and Miami-Dade, respectively. *Culex quinquefasciatus* was the second most abundant species in Brownsville with 4,473 specimens; however, it was only collected from 6 PM to 7 AM. On the other hand, *Cx. quinquefasciatus* was not abundant in Miami-Dade, it was the fifth most abundant species collected (237 specimens), mostly between dusk and dawn (Tables S1 and S2).

### Species richness

Species richness varied greatly according to the time of the day with most species being more common between dusk and dawn. In Brownsville, the highest number of species was 15 at 10 PM and the lowest was 3 at 11 AM, 1 PM, and 2 PM. In Miami-Dade, the highest number of species collected was 14 at 9 PM and the lowest was 4 species at 9 AM (Fig. 1, Fig. S1).

**Figure 1.**
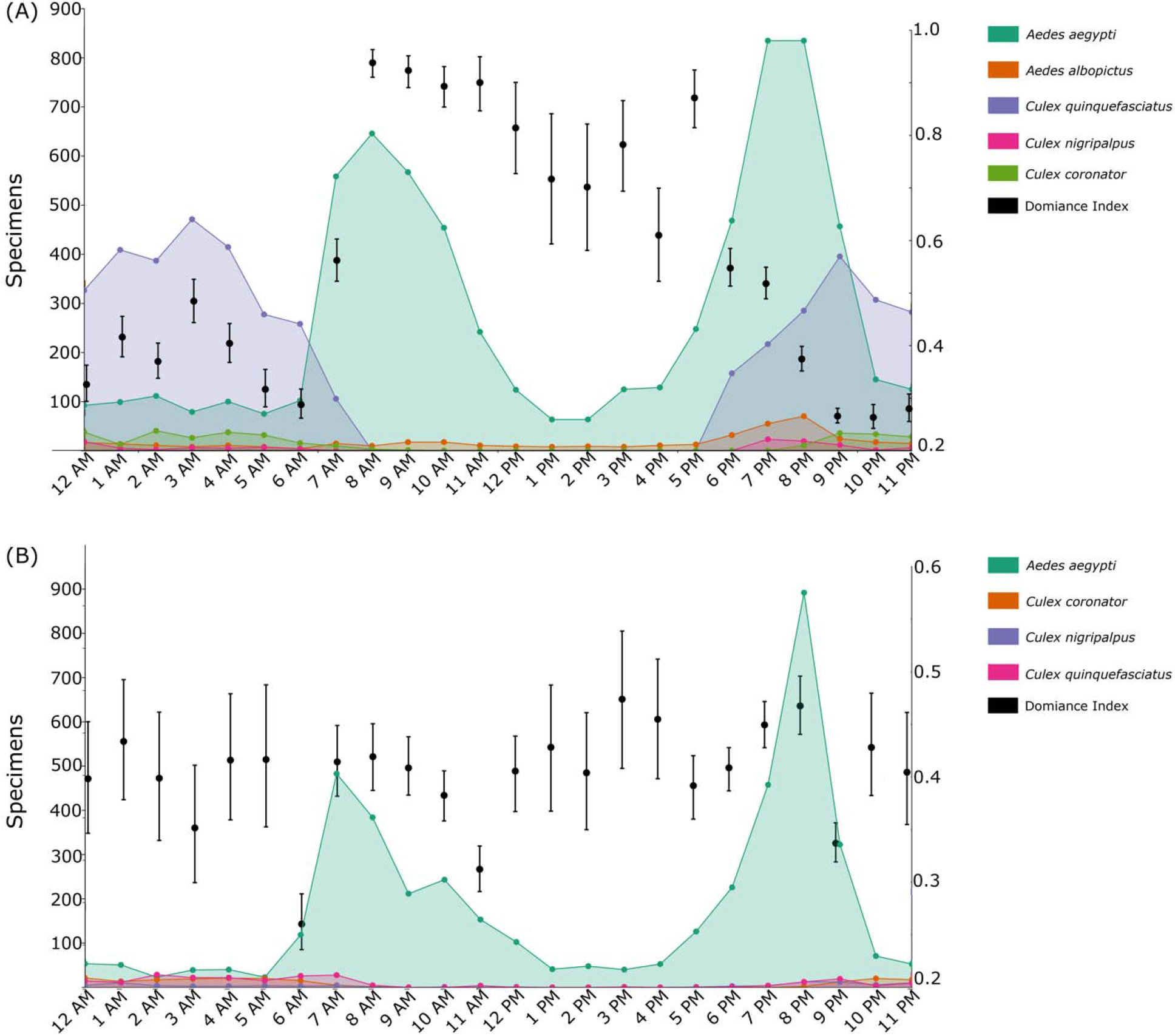
Diel activity patterns of mosquito populations collected from May to November 2019 in (A) Brownsville, Texas; and (B) Miami-Dade, Florida. The figure reports vector mosquito species only; Figure S1 shows all the collected species. The Dominance Index is shown in black (mean and 95% CI).

The PERMANOVA analyses using the Bray-Curtis and Jaccard indices yielded significant results for the diel activity patterns of mosquito species in both Brownsville and Miami-Dade. The subsequent SIMPER analyses resulted in distinct scenarios. In Brownsville, *Psorophora cyanescens, Culex erraticus, Aedes sollicitans*, and *Psorophora ciliata* contributed approximately 75% of all the variation in the mosquito community composition. Important mosquito vector species such as *Cx. nigripalpus* (7.6%), *Cx. quinquefasciatus* (<0.001%), *Ae. aegypti* (<0.001%), and *Cx. coronator* (<0.001%) did not contribute substantially to the variation (Table 1). A different scenario was observed in Miami-Dade, with *Ae. aegypti*, *Aedes taeniorhynchus*, and *Wyeomyia vanduzeei* accounting for more than 80% of all the variation in the mosquito community composition. Vector species such as *Ae. albopictus*, *Cx. quinquefasciatus*, *Cx. coronator*, and *Cx. nigripalpus* did not contribute substantially to the variation in the community composition (Tables S3 and S4).

**Table 1.**
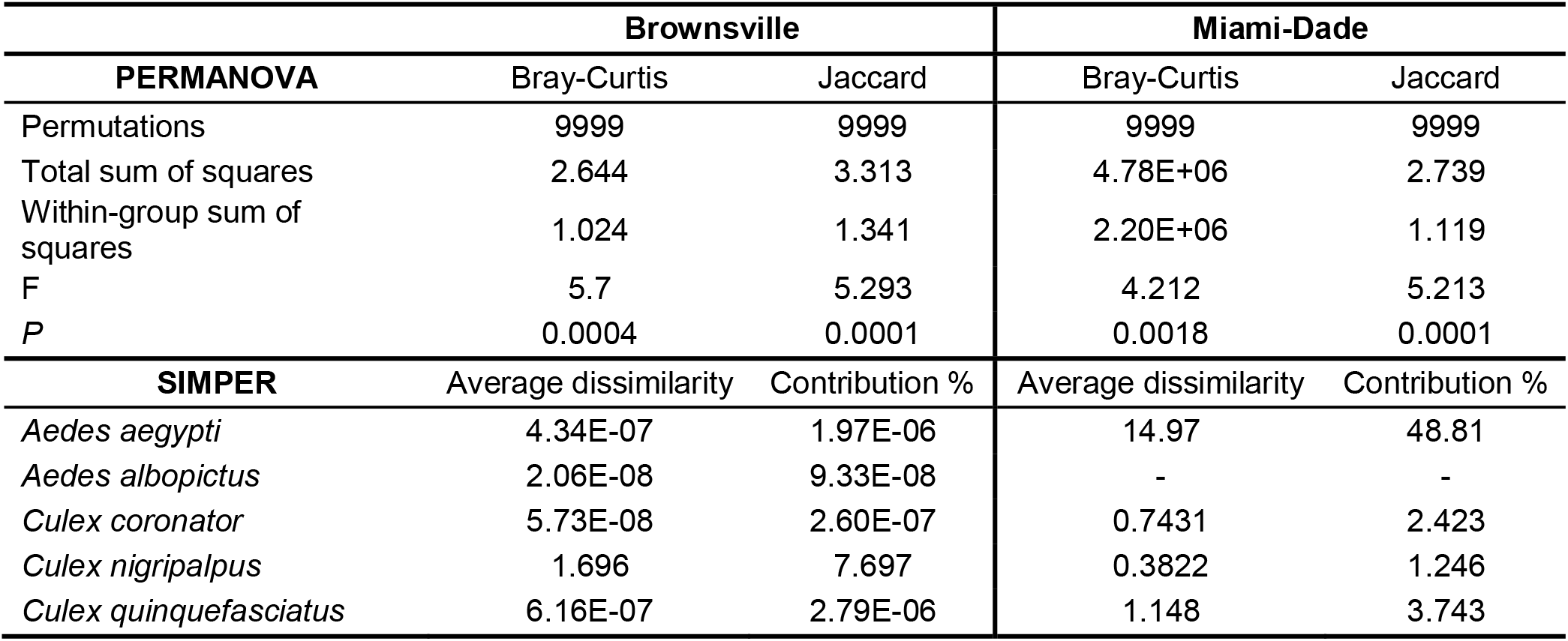
Permutational Multivariate Analysis of Variance (PERMANOVA) and SIMPER (Similarity Percentage) analysis of which species contributed the most to the observed differences in Brownsville, Texas and Miami-Dade, Florida.

### Effectiveness of adulticide spraying at different hours of the day

We leveraged a model of the mosquito population dynamics to test the effectiveness of adulticide interventions at different times of the day (Fig. S2). Specifically, we considered two scenarios representing two distinct epidemiological situations. Scenario one is during an ongoing arboviral outbreak in which the mosquitoes vector populations must be reduced quickly to curtail the epidemic. Scenario two is an effort to decrease the likelihood of an arboviral outbreak by reducing the sizes of the mosquito vector populations. In scenario two, adulticide applications may be especially important in low-resource settings or in specific situations such as before outdoor sports events or during holidays in touristic areas when source reduction or the application of larvicides are not practical. Therefore, two different adulticide application regimes to reduce adult mosquito population sizes were evaluated; (i) adulticide applications twice a day for five consecutive days and (ii) adulticide applications once a week for two months. For sensitivity analysis, we considered two alternative frequencies, once a month for two months and once every other week for two months. The main analysis considers the adulticide to be 50% effective; 30% and 70% are used for sensitivity analyses presented in the Supplementary Material (Fig. S3 and S4).

The estimated effectiveness of adulticide spraying to control populations of adult vector mosquito species varied greatly according to the hour of the intervention for both daily and weekly applications. Regarding the spraying of adulticide twice per day for 5 consecutive days, even though *Ae. aegypti* was collected during all 24 hours of the day it was more abundant in the morning and evening and the maximum effectiveness was obtained between 8 AM and 12 PM and 7 PM and 11 PM in Brownsville and Miami-Dade, respectively. Our results show that when the adulticide interventions were done during these hours of the day reaching more than 80% effectiveness in reducing *Ae. aegypti* populations. The results for *Ae. albopictus* indicate that adulticide interventions in the evening (between 5 PM and 11 PM) were the most effective reaching a maximum 95.9% reduction (95% CI: 95.3 to 96.4%). Similar results were found for *Cx. quinquefasciatus*, *Cx. coronator*, and *Cx. nigripalpus* in which adulticide applications during the day from 9 AM to 6 PM were mostly ineffective, having a negligible impact on the abundance of mosquitoes from this species. On the other hand, adulticide applications between 6 PM and 8 AM achieved the best outcomes with a maximum of 95.7% reduction (95% CI: 95.0 to 95.9%) in the abundance of these species (Fig. 2A and 2B).

**Figure 2.**
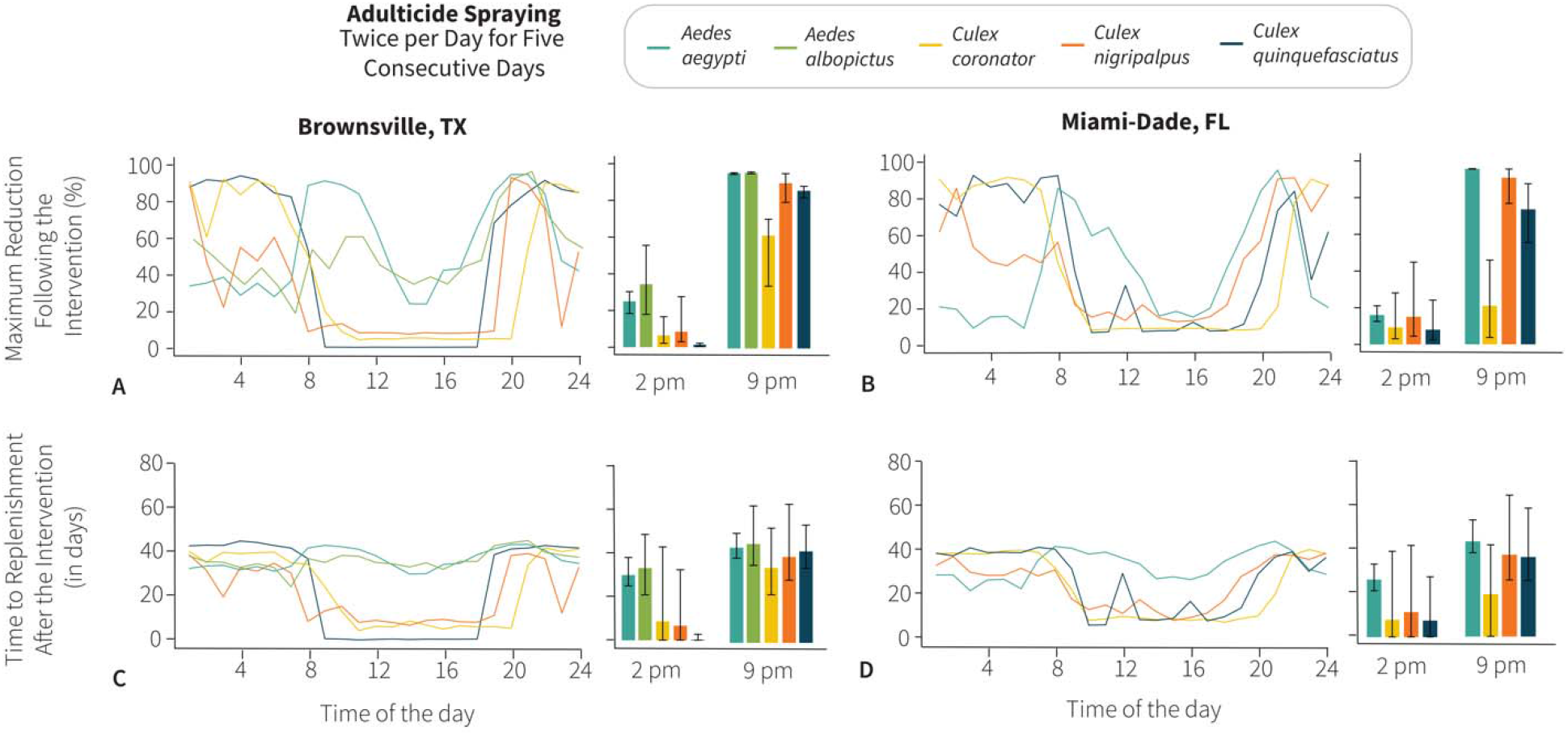
Effectiveness of adulticide spraying to control populations of adult vector mosquito species at different hours of the day. Percentage reduction of populations of vector mosquito species following adulticide application at different hours of the day twice per day for five consecutive days in (A) Brownsville, Texas and (B) Miami-Dade, Florida; Bar plots show the maximum reduction of interventions at 2 PM and 9 PM with a 95% confidence interval. Number of days after adulticide intervention at different hours of the day twice per day for five consecutive days until the mosquito populations reached the abundance levels prior to the intervention in (C) Brownsville, Texas and (D) Miami-Dade, Florida. Bar plots show the result of interventions at 2 PM and 9 PM with a 95% confidence interval for the time needed for mosquitoes to reach the abundance levels prior to the adulticide intervention.

Furthermore, the time of the adulticide intervention was associated with the number of days it took for a mosquito population to return to abundance levels prior to the chemical intervention. It would take 43.8 (95% CI: 38.9 to 50.4) days for the *Ae. aegypti* population to return to abundance levels prior to the adulticide intervention at its peak activity hours, and adulticide interventions during hours *Ae. aegypti* is not active lead to a less effective reduction, taking 30.2 (95% CI: 25.2 to 38.3) days to return to the abundance levels prior to the chemical intervention.

Adulticide interventions between 6 PM to 8 AM were the most effective in substantially reducing *Cx. coronator, Cx. nigripalpus*, and *Cx. quinquefasciatus* mosquitoes. As a result, it would take 42.1 (95% CI: 34.3 to 54.2) days for the *Cx. quinquefasciatus* population to reach the same level prior to the adulticide intervention, 34.5 (95% CI: 22.1 to 52.8) days for *Cx. coronator*, and 39.5 (95% CI: 28.8 to 63.7) days for *Cx. nigripalpus*. However, adulticide interventions were much less effective and often generated negligible results in reducing *Culex* populations if carried out between 8 AM and 6 PM, especially in Brownsville (Fig. 2C and 2D, Fig. S5).

The results for adulticide application once per week for two months indicate that mosquito populations can be reduced by up to 40.7% when the intervention is carried out when mosquitoes are more active. The *Ae. aegypti* population from Brownsville was substantially reduced when the adulticide application was carried out between 8 AM and 12 PM and reached optimum results at 8 PM and 9 PM, whereas in Miami-Dade adulticide applications in the morning were not as effective as in Brownsville and the most reduction of 39.8% (95% CI: 39.6 to 40.1%) was obtained in the evening at 9 PM. The results for *Ae. albopictus* in Brownsville were similar to those for *Ae. aegypti* in Miami-Dade. The most effective hour to control *Ae. albopictus* populations was between 7 PM and 10 PM, reaching a 40.7% (95% CI: 39.6 to 42.0%) reduction at 9 PM. Adulticide application at any other hour of the day resulted in a maximum of 15.9% reduction (95% CI: 9.9 to 22.8%).

The results for *Cx. quinquefasciatus* and *Cx. coronator* were similar in both Brownsville and Miami-Dade. In Brownsville the maximum reduction was between 7 PM and 8 AM (36.7% reduction at 4 AM; 95% CI: 36.2 to 37.3%) for *Cx. quinquefasciatus* and 9 PM and 7 AM (33.2% reduction at 3 AM; 95% CI: 23.7 to 37.6%) for *Cx. coronator*. In Miami-Dade the maximum reduction was between 9 PM and 8 AM (31.7% at 5 AM; 95% CI: 20.2 to 38.1%) for *Cx. coronator* and between 9 PM and 9 AM (33.3% reduction with 95% CI: 22.6 to 37.8% at 3 AM; 31.1% reduction with 95% CI: 20.3 to 37.5% at 7 AM) for *Cx. quinquefasciatus*. The most effective hour of the day to control *Cx. nigripalpus* was found to be between 9 PM to 11 PM (29.9% reduction at 9 PM; 95% CI: 18.9 to 37.6%) and between 12 AM to 2 AM in Brownsville. In Miami-Dade the most effective hour of the day to control *Cx. nigripalpus* was between 8 PM and 2 AM (31.0% reduction with 95% CI: 15.4 to 38.5%) and at 9 PM (31.9% reduction with 95% CI: 19.4 to 39.1%) (Fig. 3, Fig. S6).

**Figure 3.**
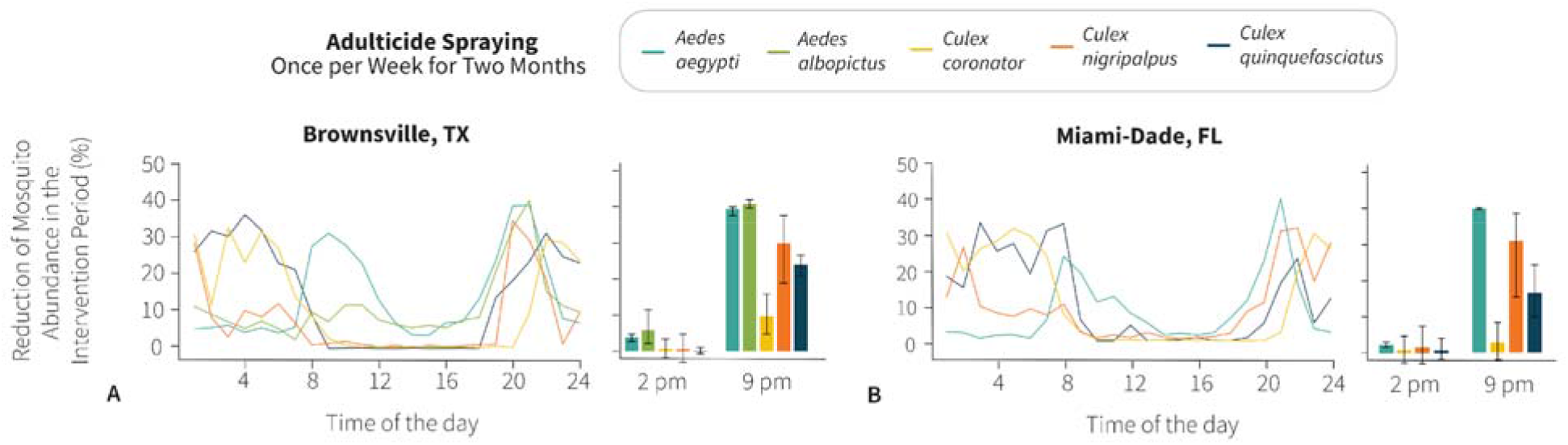
Average reduction in the abundance of populations of vector mosquito species during adulticide interventions comprised of weekly adulticide spraying for two months at different hours of the day in (E) Brownsville, Texas; and (F) Miami-Dade, Florida. Bar plots show the average reduction in the abundance of populations of vector mosquito species after adulticide interventions at 2 PM and 9 PM with a 95% confidence interval.

In terms of timing of adulticide application, results assuming a different frequency of the intervention and those assuming a different efficacy of the adulticide are consistent with those reported in the main analysis, albeit with lower absolute reductions (Fig. S3 and S4).

## Discussion

Scientific evidence on data on diel activity patterns of different vector mosquito species in different areas is lacking. This is because diel activity patterns are not routinely investigated during the development and the implementation of mosquito control strategies. The current mosquito control strategies try to target peak activity times for adulticide applications, but rarely account for local and geographic variations in activity patterns of the target vector populations, thus, failing to reach the maximum effectiveness in controlling adult populations of the vector species. Our results showed that the mosquito community composition and abundance varied significantly throughout the day in both Brownsville, Texas and Miami-Dade, Florida. However, in Brownsville, epidemiologically important mosquito species did not significantly contribute to the hourly variation in the mosquito community composition, whereas in Miami-Dade, *Ae. aegypti*, was the most dominant species and accounted for approximately 50% of the variation in the mosquito community composition. Even though fewer species were collected in Brownsville and *Ae. aegypti* was the dominant species from 8 AM to 5 PM, other epidemiologically relevant mosquito species (e.g., *Cx. quinquefasciatus* and *Ae. albopictus*) were abundantly found between dusk and dawn. On the other hand, *Ae. aegypti* was the most abundant species collected in Miami-Dade at all hours of the day, and despite the higher species richness when compared to Brownsville, the bimodal diel activity pattern of *Ae. aegypti* was a major driver for the variation found in the mosquito community composition.

Considering the natural variations in the diel activity patterns of mosquito vector species present in Brownsville and Miami-Dade, we modeled adulticide applications at every hour of the day. Our results indicate that the effectiveness of adulticide applications varied greatly according to the hour of the day and most effective at the peak activity time of the target mosquito vector species. Should the intervention be aimed at reducing mosquito populations as much as possible in the shortest amount of time, as would be the case when an arbovirus outbreak is ongoing, we simulated a scenario where the adulticide is sprayed twice per day for 5 consecutive days. In the case of *Ae. aegypti* (the primary vector of Zika, dengue, and chikungunya viruses in the study locations) in both Brownsville and Miami-Dade, targeting the adulticide application when they are most active, 8 AM and 8 PM, resulted in the reduction of approximately 90% of the population. However, adulticide applications targeting *Ae. aegypti* in the early morning (between 1 AM and 7 AM) reduced only around 40% of the population. By targeting *Ae. aegypti* at its peak activity hour, it would take approximately 60 days for the population to reach the same abundance as prior to the adulticide intervention. However, targeting *Ae. aegypti* in the early morning would result in a less effective mosquito control intervention with the population reaching the same numbers as prior to the adulticide intervention after 40 days. Our results show that a simple change in the adulticide spraying from 7 AM to 8 AM would substantially impact the effectiveness of the mosquito control intervention. Spraying adulticide twice per day for 5 consecutive days yielded even more pronounced results for *Cx. coronator, Cx. nigripalpus*, and *Cx. quinquefasciatus*. Our results show that spraying adulticide between 9 AM and 6 PM achieved a negligible reduction in the *Culex* species. On the other hand, adulticide spraying at 9 PM led to a 90% reduction in the number of mosquitoes. As a result, just one hour difference in targeting *Cx. quinquefasciatus* can lead to a 70% reduction at 7 PM as opposed to a negligible reduction at 6 PM. Similar results were found for *Cx. coronator* and *Cx. nigripalpus*, in which spraying at 10 PM led to a 90% reduction but at 8 PM the reduction would have been negligible. By targeting the *Culex* species at their peak activity time, their populations would need approximately 50 days to rebound to the levels prior to the adulticide spraying in contrast with no reduction if the intervention would have been done just a few hours earlier. From an epidemiological standpoint, it is important to note that a 90% reduction in the vector population would have a substantial effect on the control of an arbovirus outbreak, causing reproduction numbers of up to 10 to fall below the epidemic threshold of 1. In fact, the reproduction number (i.e., the number of secondary infections generated by an infective host through vector bites in a fully susceptible host population) for a mosquito-borne infectious disease is directly proportional to the number of adult female vectors.

When interventions are aimed at keeping mosquito vector populations to acceptable levels, for instance, to decrease the likelihood of an arbovirus outbreak, we simulated spraying adulticide in more sparse intervals for two months (e.g., once per week, once every other week, and once per month). Also, in this case, the effectiveness of the intervention was positively associated with the peak activity time of each target mosquito vector species. For instance, for *Ae. aegypti*, the estimated reduction ranged from 20.1% (95% CI: 19.7 to 20.6%) and 20.1% (95% CI: 19.5 to 20.4%) when spraying is performed at 9 PM in both Miami-Dade and Brownsville to 4.3% (95% CI: 3.1 to 5.3%) and 2.8% (95% CI: 1.9 to 3.8%) when spraying is performed at 2 PM.

Our modeling analysis considers the worst-case scenario in which the temperature is kept constant at 25 degrees Celsius – an optimal temperature for mosquito development and proliferation [24]. As such, our estimates can be considered as a lower bound of the effectiveness of the intervention. A limitation of our analysis is due to the unavailability of the development and mortality ratios for *Cx. coronator* and *Cx. nigripalpus* in the scientific literature, we have used the parameters that were available for *Cx. quinquefasciatus*. The mosquito species of the *Culex* genus are organized into 26 subgenera, and both *Cx. coronator* and *Cx. nigripalpus* share the same subgeneric classification as *Cx. quinquefasciatus*; they all belong to the subgenus *Culex* [25]. Therefore, even though these are different species they share similar biological traits and the parameters of *Cx. quinquefasciatus* are reasonable approximations for *Cx. coronator* and *Cx. nigripalpus* (Table S5). Another limitation is that the model considers a single mosquito population at the time, thus ignoring possible competition between species. However, we do not expect the competition for resources to be an important driver for the proliferation of the species selected for this study as they exploit different resources in the urban environment and have distinct population dynamics [26,27].

Adulticide spraying during the day would potentially increase the risk of exposure of the general public to adulticides and should be considered in the development of mosquito control strategies. Furthermore, different propellant mechanisms (e.g., Buffalo Turbine, Grizzly ULV Sprayer, airplane, etc.) may have different levels of effectiveness due to intrinsic limitations in their operation mechanisms, droplet size, as well as due to different weather conditions (e.g., air temperature and wind speed), presence of vegetation, and other factors present in the target area.

Despite being one of the most important aspects for the development of effective mosquito control strategies based on adulticide spraying and assessing the exposure risk to infective bites, the only recent studies focusing on the diel activity patterns of *Ae. aegypti* in the contiguous United States were done in St. Augustine, Florida in 2017 by Smith et. al. [28] and in 2020 in Miami-Dade, Florida and Brownsville, Texas by Mutebi et al. [29]. Both studies show a clear bimodal pattern of peak activity of *Ae. aegypti* in the morning and evening.

Adulticides are commonly used in the United States during emergency situations to control and prevent arbovirus outbreaks. For instance, they were widely used during the Zika virus outbreak in Miami-Dade, Florida [3,30,31]. Our results have great implications to improve the effectiveness of mosquito control strategies. Indeed, diel activity patterns of vector mosquito species should be considered when developing future mosquito control strategies.

In case of an arbovirus outbreak, in addition to adulticide application, it is common practice, as well as recommended by CDC guidelines for surveillance, prevention, and control of West Nile Virus [32,33], to establish a perimeter and remove potential aquatic habitats, spray larvicide and adulticide at the point source location of the infection and patient’s home address in response to imported and locally transmitted arboviral infections [34]. The use of adulticide disregarding the diel activity of the target mosquito species can lead not only to ineffective control interventions but, more importantly, the indiscriminate use of adulticide, even during emergency situations, can have deleterious results by increasing the levels of adulticide resistance of the target mosquito population which may lead to decreased effectiveness of adulticide(s) [35]. Targeting adult mosquito vector species with adulticide spraying-based control strategies during their peak activity will not only result in more effective control strategies but will also result in the need to spray adulticides fewer times, potentially reducing the development of adulticide resistance by the target mosquito species and the exposure of non-target species.

## Conclusion

This study serves as a cornerstone for future studies to improve the effectiveness of mosquito control strategies that employ adulticide applications. Our results point to a well-defined diel activity pattern in the peak activity of mosquito species reaching maximum relative abundance at specific times in both Brownsville, Texas and Miami-Dade, Florida. Furthermore, the effectiveness of adulticide applications varied greatly according to the hour of the day with 9 PM being the most effective time of the day to control populations of all vector mosquito species in Miami-Dade and Brownsville. Our results indicate that the timing of adulticide spraying interventions should be carefully considered by local authorities based on the ecology of mosquito species in the focus area.

## Methods

### Mosquito collection and study sites

Mosquitoes were collected by BG-Sentinel 2 traps (Biogents AG, Regensburg, Germany) baited with BG Lures and dry ice as a source of carbon dioxide in Miami-Dade County, Florida and Brownsville, Texas as described in Mutebi et al. [29]. The study locations were chosen due to their history of arboviral outbreaks, including West Nile, yellow fever, dengue, and Zika viruses. In each city, four collection sites were selected to capture the natural variation in the diel activity patterns of mosquito species due to their inherent socio-ecological features. BG-Sentinel 2 traps are the gold standard to assess the community composition of mosquitoes in urban environments. They mimic a potential host by using BG-Lures that release attractants consisting of carbon dioxide, lactic acid, ammonia, and caproic acid, substances that are found on animal skin thus attracting female mosquitoes seeking blood sources. In addition, the BG traps were baited with dry ice to release carbon dioxide, which has been shown to significantly increase the capture of diverse mosquito species [36]. Male mosquitoes collected during this study were considered accidental catches and, since they were not consistently collected, they were not considered informative and were not included in the analyses.

Mosquito collections were made from May 2019 to November 2019 as previously described by Mutebi et. al. [29]. Each BG-Sentinel trap was monitored every hour for 96 hours at each location once every month from May to November 2019. Every hour, the BG-Sentinel traps were serviced by project personnel, and the collection bags were removed from the traps and replaced with empty ones. The bags of mosquitoes were then labeled and transported to the laboratory under refrigeration, where they were subsequently identified to species on chill tables by using entomological keys [37].

### Diversity Analysis

To assess variations in the diel activity patterns of mosquito species in each city we organized the data into six groups (group 1 = 12 AM to 3 AM; group 2 = 4 AM to 7 AM; group 3 = 8 AM to 11 AM; group 4 = 12 PM to 3 PM; group 5 = 4 PM to 7 PM; and group 6 = 8 PM to 11 PM) and performed a PERMANOVA with 9,999 permutations based on Bray-Curtis (abundance-based) and the Jaccard (incidence-based) indices [38]. Then we used the SIMPER method for assessing which species has contributed the most to the observed differences between groups of samples [39]. To assess hourly variations in the mosquito community composition, we used the Dominance (1-Simpson) index, in which values closer to 0 indicate species are equally present and values closer to 1 indicate the presence of highly dominant species. Analyses were done using PAST v4 [40].

### Stochastic Compartmental Model

We used a stochastic compartmental model to simulate the population dynamics of mosquito species to determine the effectiveness of adulticide applications at different times of the day. We selected the most epidemiologically important mosquito vector species that were collected in Brownsville, Texas: *Aedes aegypti, Aedes albopictus, Culex coronator, Culex nigripalpus*, and *Culex quinquefasciatus*; and in Miami-Dade, Florida: *Ae. aegypti, Cx. coronator, Cx. nigripalpus*, and *Cx. quinquefasciatus*.

The mathematical model for the dynamics of the vector mosquito populations considers four developmental stages: eggs (E), larvae (L), pupae (P), and the adult female (A) population.

In our mathematical model, we assume that under suitable conditions, eggs hatch into larvae at rate d_E_ and some suffer natural mortality at rate m_E_. Upon feeding on micro-organisms, and after moulting three times, the larvae then develop into pupae at rate d_L_. In our model, we consider the carrying capacity K of the breeding sites to limit the maximum number of eggs that can be allowed to develop. As such, we assume no further competition for food among the larvae; hence they only suffer natural mortality at rate m_L_. Pupae develop to become adult mosquitoes at rate d_P_ and they die at rate m_P_. We considered a 1:1 sex ratio among adult mosquitoes [41]. In the adult stage, we consider female adult mosquitoes only as males are not epidemiologically relevant. We considered the gonotrophic cycles to occur at rate dA and nE represents the average numbers of eggs laid per each oviposition. Finally, adult mosquitoes die at rate mA. All model parameters are species-dependent.

The described process can mathematically be represented by the following system of equations:

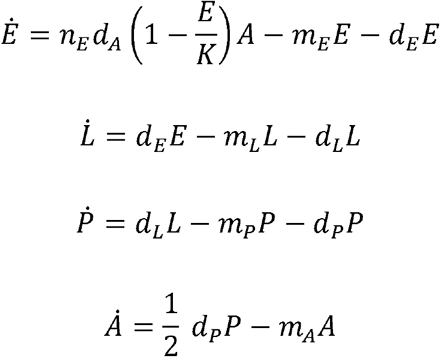

In our mathematical model, transitioning from one life stage to another is a stochastic process modelled as binomial transitions. We utilize a discrete-time stochastic version of our mathematical model, with time a step of 0.1 days. The model was implemented in R.

### Development and mortality rates

All the development and mortality rates used in our model are temperature- and species-dependent and we used a constant temperature of 25 degrees Celsius. The development and mortality rates for *Ae. albopictus* were obtained from Delatte et al. [41] and implemented as in Poletti et al. [38][30]. The larvae and pupae development ratios for *Ae. aegypti* and *Cx. quinquefasciatus* were obtained from Grech et al. [24]. The egg hatching and the gonotrophic times for the *Ae. aegypti* were obtained from Farnesi et al. [42] and Garcia-Rejon et al. [43] respectively. Additionally, the egg hatching and the gonotrophic cycle for the *Cx. quinquefasciatus* were taken from Handel [44] and Laporta & Sallum [45] respectively. The mortality rates for eggs, larvae, pupae, and adult *Cx. quinquefasciatus* mosquitoes were obtained from Ewing et al. [46] and Watanabe et al. [47] and Farnesi et al. [42], Yang et al. [48] and Yang et al. [49] for *Ae. aegypti*, respectively.

### Simulation model of adulticide spraying

Four frequencies of applications of adulticides at a given time of the day are considered: i) twice a day for 5 consecutive days, ii) once a month for two months, iii) once a week for two months, and iv) once every other week for two months. We consider adulticide spraying to affect only adult mosquitoes that are active at the time of spraying. As such, for each study location, hour of the day, and mosquito species, we estimated the fraction of active mosquitoes as: *r_ij_*(*t*) = *C_ij_*(*t*)/[*C_ij_*(*t*)], where *C_ij_*(*t*) represents the estimated number of active adult mosquitoes in location *i* = {*Brownsville, Miami*} of species *j*, at time of the day *t*. To account for the uncertainty in the number of captured mosquitoes at each time of the day, *C_ij_*(*t*) has been sampled from a Poisson distribution of mean 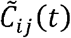, representing the observed number of captured mosquitos in location *i*, of species *j*, at time of the day t. Upon spraying, the adult mosquito population is instantly reduced by *σr_ij_*(*t*), where *σ* represents the spraying efficacy. Spraying efficacy is set at 50% in the main analysis, while 30% and 70% are investigated in the supplementary analyses. To estimate the uncertainty in the effectiveness of the interventions, for each location, mosquito vector species, and time of the adulticide spraying, we performed 1,000 simulations, one for each sampled value of *C_ij_*(*t*).

## Disclaimer

The information provided, and views expressed in this publication do not necessarily reflect the official position of the Centers for Disease Control and Prevention (CDC).

---

**Supplementary Table 1.**
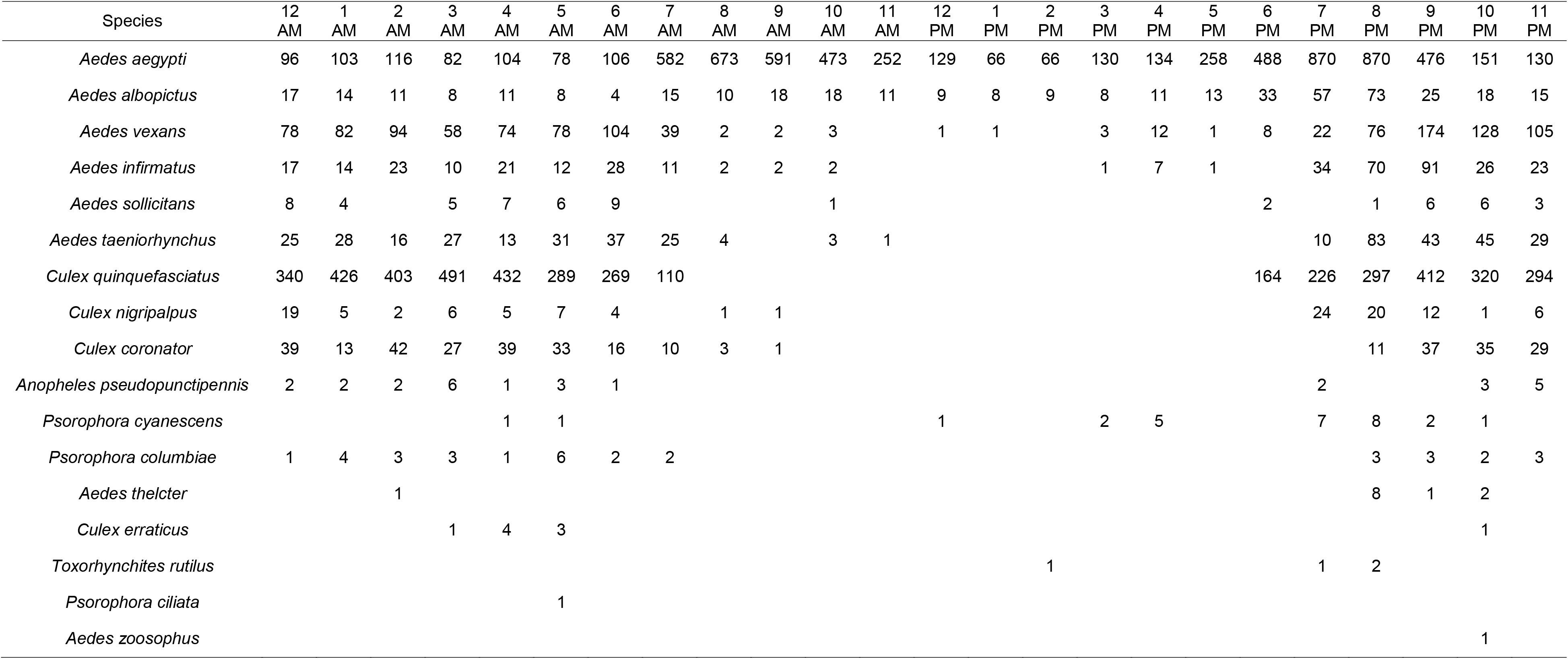
Female mosquitoes captured by BG-Sentinel 2 traps in Brownsville, Texas.

| Species | 12 AM | 1 AM | 2 AM | 3 AM | 4 AM | 5 AM | 6 AM | 7 AM | 8 AM | 9 AM | 10 AM | 11 AM | 12 PM | 1 PM | 2 PM | 3 PM | 4 PM | 5 PM | 6 PM | 7 PM | 8 PM | 9 PM | 10 PM | 11 PM |
| --- | --- | --- | --- | --- | --- | --- | --- | --- | --- | --- | --- | --- | --- | --- | --- | --- | --- | --- | --- | --- | --- | --- | --- | --- |
| *Aedes aegypti* | 96 | 103 | 116 | 82 | 104 | 78 | 106 | 582 | 673 | 591 | 473 | 252 | 129 | 66 | 66 | 130 | 134 | 258 | 488 | 870 | 870 | 476 | 151 | 130 |
| *Aedes albopictus* | 17 | 14 | 11 | 8 | 11 | 8 | 4 | 15 | 10 | 18 | 18 | 11 | 9 | 8 | 9 | 8 | 11 | 13 | 33 | 57 | 73 | 25 | 18 | 15 |
| *Aedes vexans* | 78 | 82 | 94 | 58 | 74 | 78 | 104 | 39 | 2 | 2 | 3 |  | 1 | 1 |  | 3 | 12 | 1 | 8 | 22 | 76 | 174 | 128 | 105 |
| *Aedes infirmatus* | 17 | 14 | 23 | 10 | 21 | 12 | 28 | 11 | 2 | 2 | 2 |  |  |  |  | 1 | 7 | 1 |  | 34 | 70 | 91 | 26 | 23 |
| *Aedes sollicitans* | 8 | 4 |  | 5 | 7 | 6 | 9 |  |  |  | 1 |  |  |  |  |  |  |  | 2 |  | 1 | 6 | 6 | 3 |
| *Aedes taeniorhynchus* | 25 | 28 | 16 | 27 | 13 | 31 | 37 | 25 | 4 |  | 3 | 1 |  |  |  |  |  |  |  | 10 | 83 | 43 | 45 | 29 |
| *Culex quinquefasciatus* | 340 | 426 | 403 | 491 | 432 | 289 | 269 | 110 |  |  |  |  |  |  |  |  |  |  | 164 | 226 | 297 | 412 | 320 | 294 |
| *Culex nigripalpus* | 19 | 5 | 2 | 6 | 5 | 7 | 4 |  | 1 | 1 |  |  |  |  |  |  |  |  |  | 24 | 20 | 12 | 1 | 6 |
| *Culex coronator* | 39 | 13 | 42 | 27 | 39 | 33 | 16 | 10 | 3 | 1 |  |  |  |  |  |  |  |  |  |  | 11 | 37 | 35 | 29 |
| *Anopheles pseudopunctipennis* | 2 | 2 | 2 | 6 | 1 | 3 | 1 |  |  |  |  |  |  |  |  |  |  |  |  | 2 |  |  | 3 | 5 |
| *Psorophora cyanescens* |  |  |  |  | 1 | 1 |  |  |  |  |  |  | 1 |  |  | 2 | 5 |  |  | 7 | 8 | 2 | 1 |  |
| *Psorophora columbiae* | 1 | 4 | 3 | 3 | 1 | 6 | 2 | 2 |  |  |  |  |  |  |  |  |  |  |  |  | 3 | 3 | 2 | 3 |
| *Aedes thelcter* |  |  | 1 |  |  |  |  |  |  |  |  |  |  |  |  |  |  |  |  |  | 8 | 1 | 2 |  |
| *Culex erraticus* |  |  |  | 1 | 4 | 3 |  |  |  |  |  |  |  |  |  |  |  |  |  |  |  |  | 1 |  |
| *Toxorhynchites rutilus* |  |  |  |  |  |  |  |  |  |  |  |  |  |  | 1 |  |  |  |  | 1 | 2 |  |  |  |
| *Psorophora ciliata* |  |  |  |  |  | 1 |  |  |  |  |  |  |  |  |  |  |  |  |  |  |  |  |  |  |
| *Aedes zoosophus* |  |  |  |  |  |  |  |  |  |  |  |  |  |  |  |  |  |  |  |  |  |  | 1 |  |

**Supplementary Table 2.**
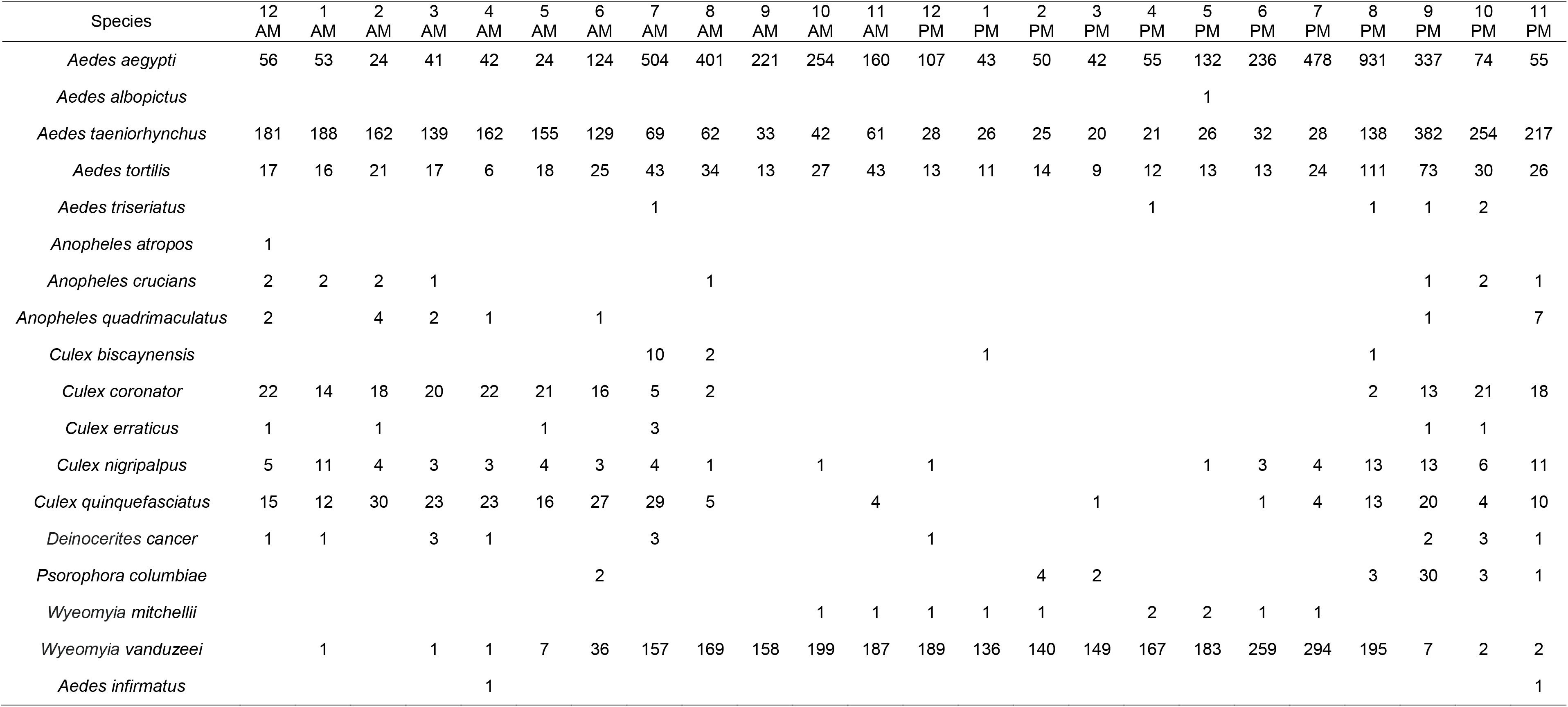
Female mosquitoes captured by BG-Sentinel 2 traps in Miami-Dade, Florida.

**Supplementary Table 3.** SIMPER (Similarity Percentage) analysis of which species contributed the most to the observed differences in Brownsville, Texas.

| Species | Average<br>dissimilarity | Contribution % | Cumulative<br>contribution % | Mean<br>1 | Mean<br>2 |
| --- | --- | --- | --- | --- | --- |
| *Psorophora cyanescens* | 5.434 | 24.65 | 24.65 | 2.15E+09 | 1.07E+09 |
| *Culex erraticus* | 4.973 | 22.56 | 47.22 | 1.61E+09 | 1.07E+09 |
| *Aedes sollicitans* | 3.12 | 14.16 | 61.37 | 5.37E+08 | 5.37E+08 |
| *Psorophora ciliata* | 2.865 | 13 | 74.37 | 2.15E+09 | 1.61E+09 |
| *Aedes thelcter* | 2.257 | 10.24 | 84.61 | 1.61E+09 | 2.15E+09 |
| *Culex nigripalpus* | 1.696 | 7.697 | 92.3 | 8 | 5.37E+08 |
| *Anopheles pseudopunctipennis* | 1.696 | 7.697 | 100 | 3 | 5.37E+08 |
| *Culex quinquefasciatus* | 6.16E-07 | 2.79E-06 | 100 | 415 | 275 |
| *Aedes aegypti* | 4.34E-07 | 1.97E-06 | 100 | 99.3 | 218 |
| *Aedes vexans* | 8.50E-08 | 3.86E-07 | 100 | 78 | 73.8 |
| *Culex coronator* | 5.73E-08 | 2.60E-07 | 100 | 30.3 | 24.5 |
| *Aedes taeniorhynchus* | 3.81E-08 | 1.73E-07 | 100 | 24 | 26.5 |
| *Aedes infirmatus* | 3.02E-08 | 1.37E-07 | 100 | 16 | 18 |
| *Aedes albopictus* | 2.06E-08 | 9.33E-08 | 100 | 12.5 | 9.5 |
| *Psorophora colombiae* | 8.65E-09 | 3.92E-08 | 100 | 2.75 | 2.75 |
| *Toxorhynchites rutilus* |  |  | 100 | 2.15E+09 | 2.15E+09 |
| *Aedes zoosophus* |  |  | 100 | 2.15E+09 | 2.15E+09 |

**Supplementary Table 4.**
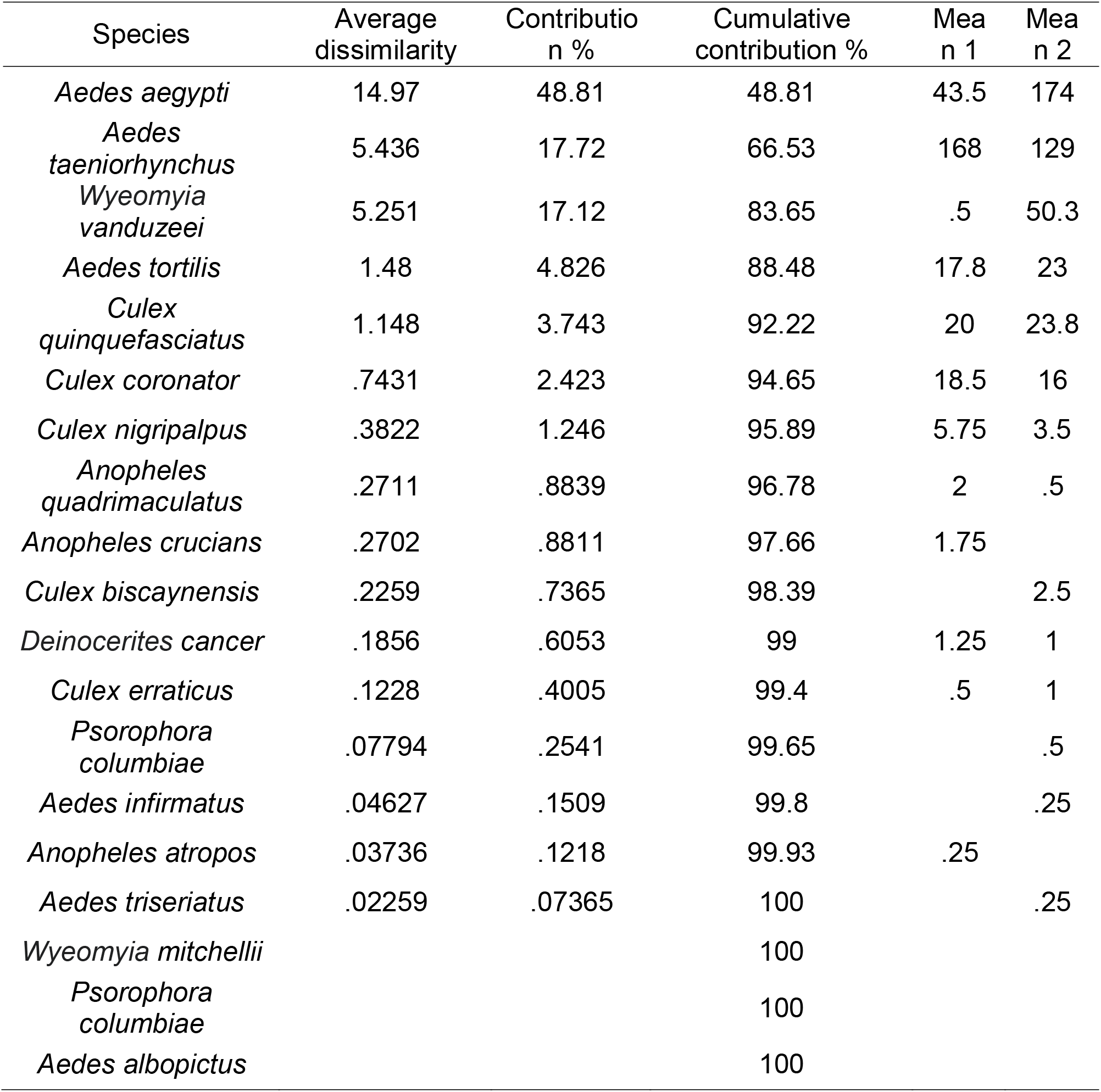
SIMPER (Similarity Percentage) analysis of which species contributed the most to the observed differences in Miami-Dade, Florida.

| Species | Average<br>dissimilarity | Contribution % | Cumulative<br>contribution % | Mean 1 | Mean 2 |
| --- | --- | --- | --- | --- | --- |
| *Aedes aegypti* | 14.97 | 48.81 | 48.81 | 43.5 | 174 |
| *Aedes taeniorhynchus* | 5.436 | 17.72 | 66.53 | 168 | 129 |
| *Wyeomyia vanduzeei* | 5.251 | 17.12 | 83.65 | .5 | 50.3 |
| *Aedes tortilis* | 1.48 | 4.826 | 88.48 | 17.8 | 23 |
| *Culex quinquefasciatus* | 1.148 | 3.743 | 92.22 | 20 | 23.8 |
| *Culex coronator* | .7431 | 2.423 | 94.65 | 18.5 | 16 |
| *Culex nigripalpus* | .3822 | 1.246 | 95.89 | 5.75 | 3.5 |
| *Anopheles quadrimaculatus* | .2711 | .8839 | 96.78 | 2 | .5 |
| *Anopheles crucians* | .2702 | .8811 | 97.66 | 1.75 |  |
| *Culex biscaynensis* | .2259 | .7365 | 98.39 |  | 2.5 |
| *Deinocerites cancer* | .1856 | .6053 | 99 | 1.25 | 1 |
| *Culex erraticus* | .1228 | .4005 | 99.4 | .5 | 1 |
| *Psorophora columbiae* | .07794 | .2541 | 99.65 |  | .5 |
| *Aedes infirmatus* | .04627 | .1509 | 99.8 |  | .25 |
| *Anopheles atropos* | .03736 | .1218 | 99.93 | .25 |  |
| *Aedes triseriatus* | .02259 | .07365 | 100 |  | .25 |
| *Wyeomyia mitchellii* |  |  | 100 |  |  |
| *Psorophora columbiae* |  |  | 100 |  |  |
| *Aedes albopictus* |  |  | 100 |  |  |

**Supplementary Table 5.** Mosquito species development and mortality ratios.

| **Parameters** | ***Aedes***<br>***aegypti*** | ***Aedes***<br>***albopictus*** | ***Culex***<br>***coronator*** | ***Culex***<br>***nigripalpus*** | ***Culex***<br>***quinquefasciatus*** |
| --- | --- | --- | --- | --- | --- |
| d_E | 0.31 | 0.16 | 0.57 | 0.57 | 0.57 |
| d_L | 0.13 | 0.12 | 0.13 | 0.13 | 0.13 |
| d_P | 0.65 | 0.48 | 0.55 | 0.55 | 0.55 |
| d_A | 0.2 | 0.20 | 0.25 | 0.25 | 0.25 |
| m_E | 0.32 | 0.16 | 0.17 | 0.17 | 0.17 |
| m_L | 0.08 | 0.02 | 0.04 | 0.04 | 0.04 |
| m_P | 0.06 | 0.02 | 0.01 | 0.01 | 0.01 |
| m_A | 0.03 | 0.03 | 0.03 | 0.03 | 0.03 |
| n_E | 100 | 100 | 150-200 | 150-200 | 150-200 |

**Supplementary Figure 1.**
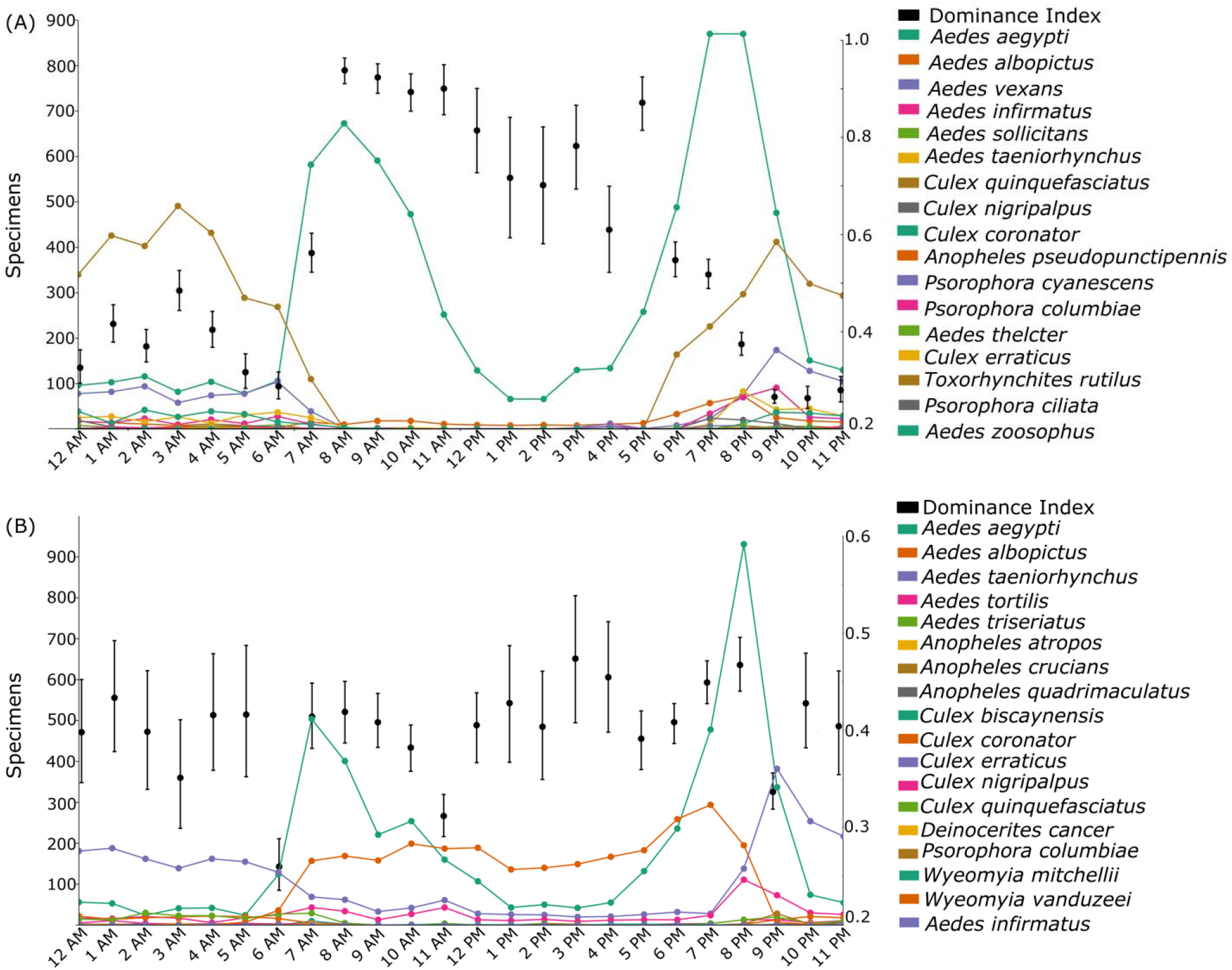
Diel activity patterns of mosquito populations collected from May to November 2019 in (A) Brownsville, Texas; and (B) Miami-Dade, Florida. Dominance Index is shown in black.

**Supplementary Figure 2.**
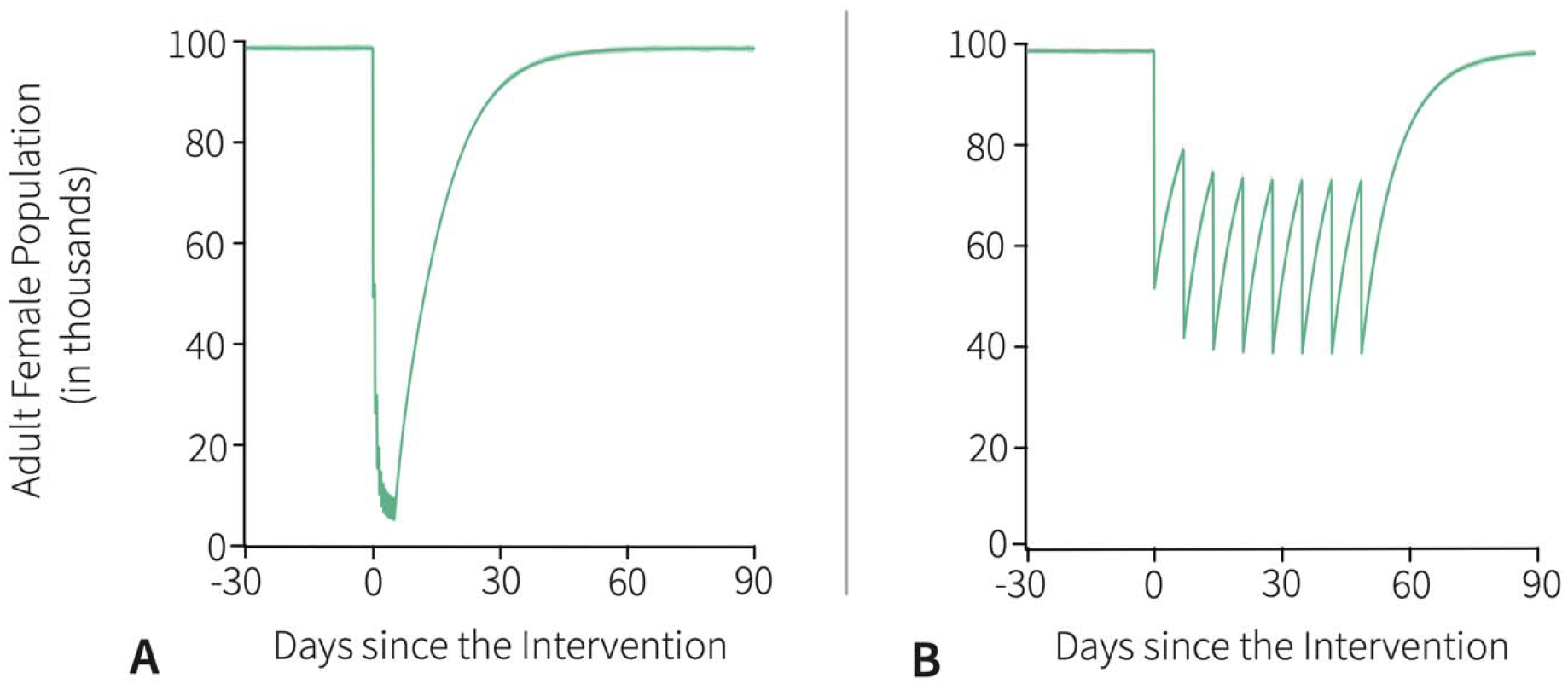
Baseline simulations and reference adulticide interventions. (A) Twice per day for five consecutive days; and (B) once per week for two months.

**Supplementary Figure 3.**
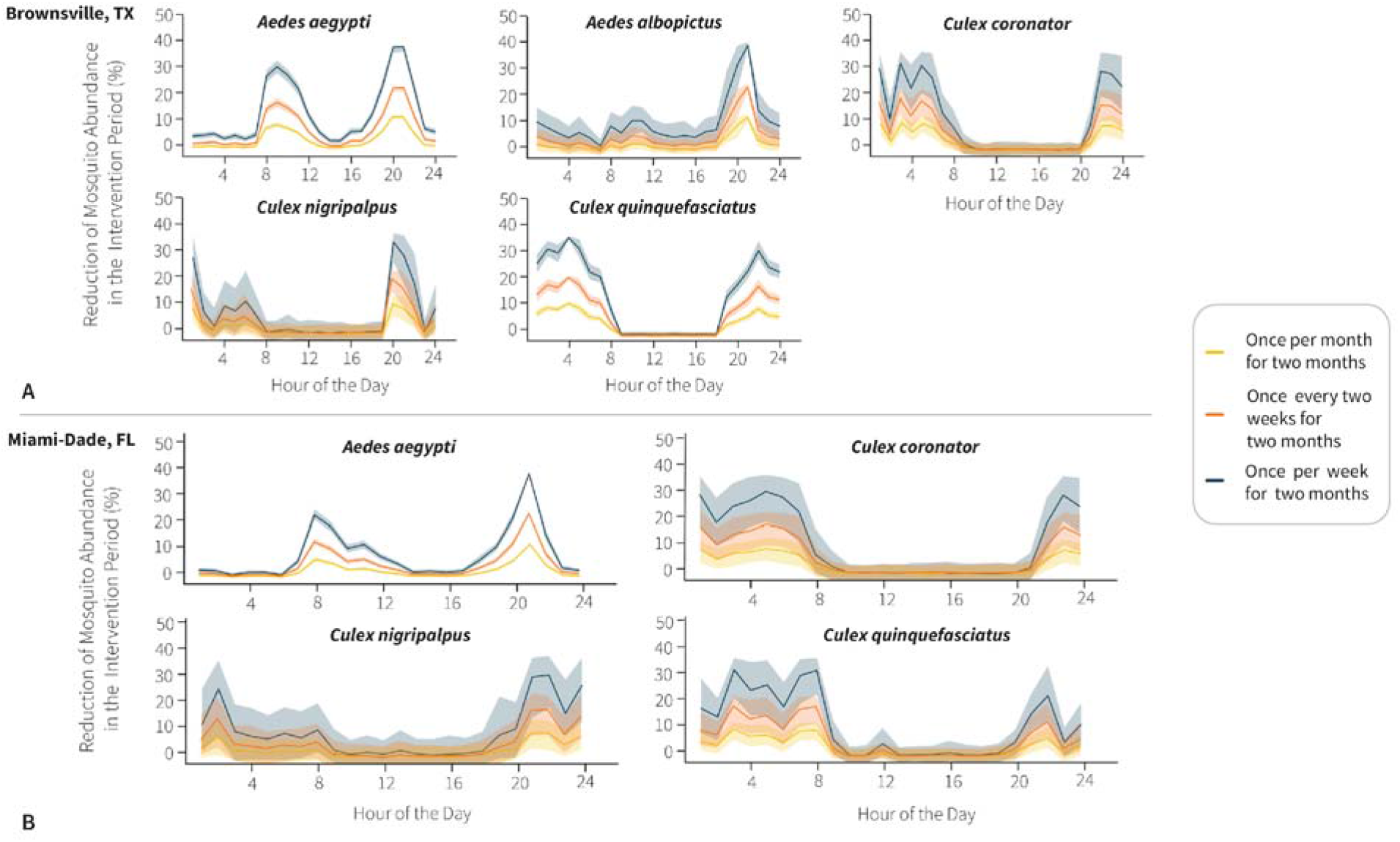
Effectiveness of different adulticide spraying strategies to control populations of adult vector mosquito species at different hours of the day. Percentage reduction of populations of vector mosquito species following different adulticide strategies at different hours of the day in (A) Brownsville, Texas and (B) Miami-Dade, Florida; Solid line = average number of mosquitoes; Light color highlights = 95% Confidence Interval.

**Supplementary Figure 4.**
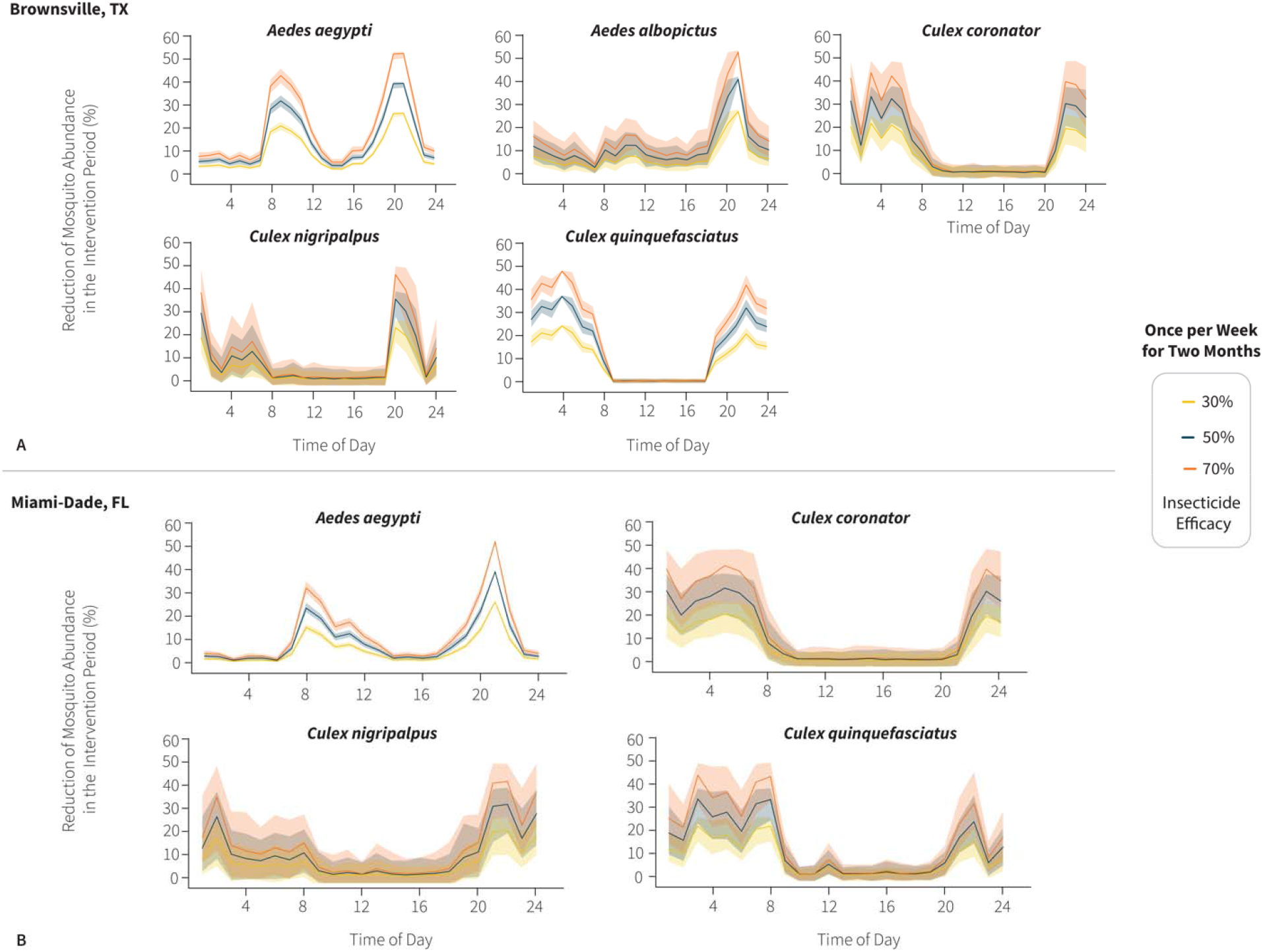
Impact of different adulticide spraying efficacy in controlling populations of adult vector mosquito species at different hours of the day. Percentage reduction of populations of vector mosquito species following different adulticide efficacies at different hours of the day in (A) Brownsville, Texas and (B) Miami-Dade, Florida; Solid line = average number of mosquitoes; Light color highlights = 95% Confidence Interval.

**Supplementary Figure 5.**
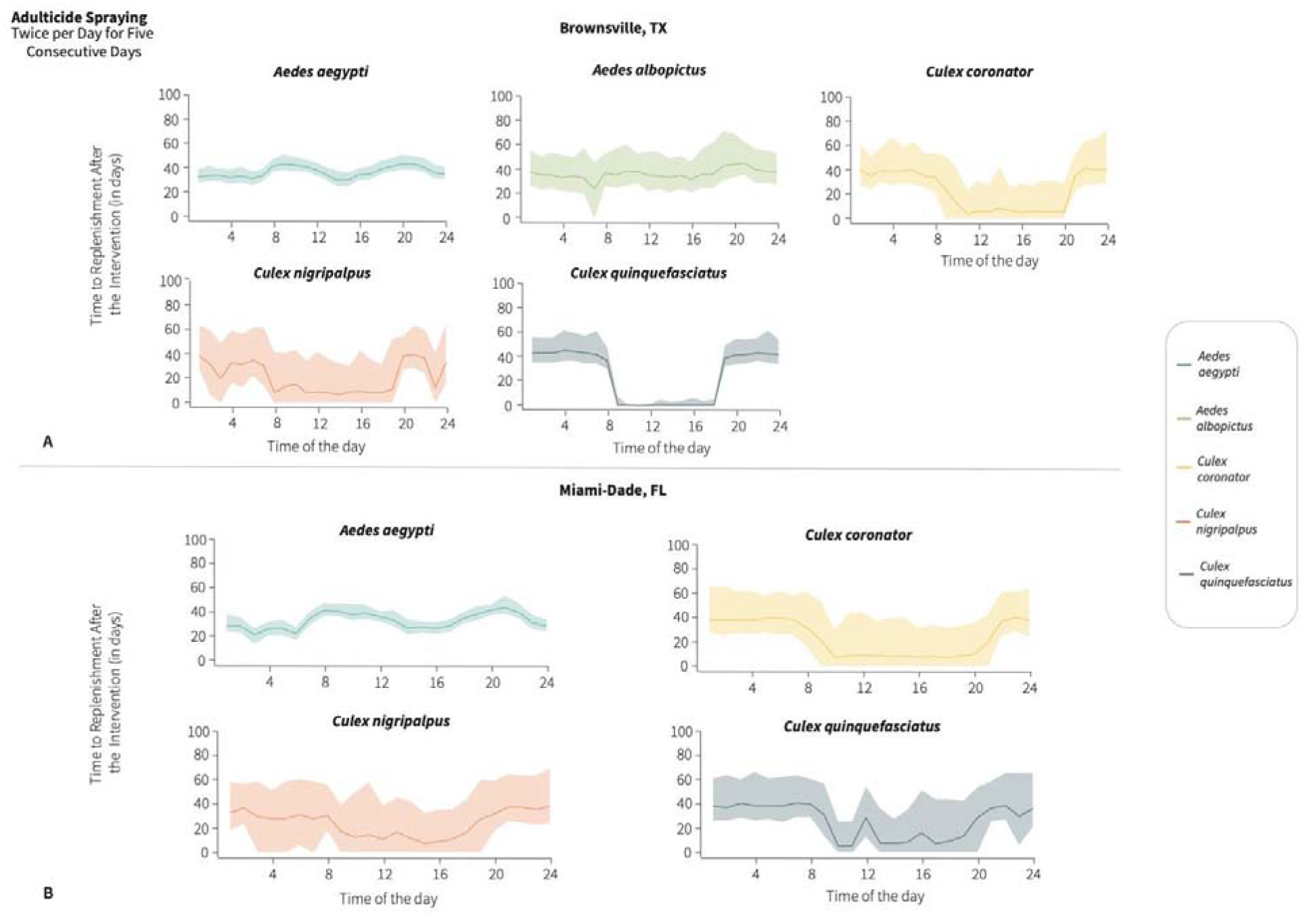
Effectiveness of adulticide spraying to control populations of adult vector mosquito species at different hours of the day. Percentage reduction of populations of vector mosquito species following adulticide application at different hours of the day twice per day for five consecutive days in (A) Brownsville, Texas and (B) Miami-Dade, Florida; Number of days after adulticide intervention at different hours of the day twice per day for five consecutive days until the mosquito populations reached the abundance levels prior to the intervention in (C) Brownsville, Texas and (D) Miami-Dade, Florida. Solid line = average number of mosquitoes; Light color highlights = 95% Confidence Interval.

**Supplementary Figure 6.**
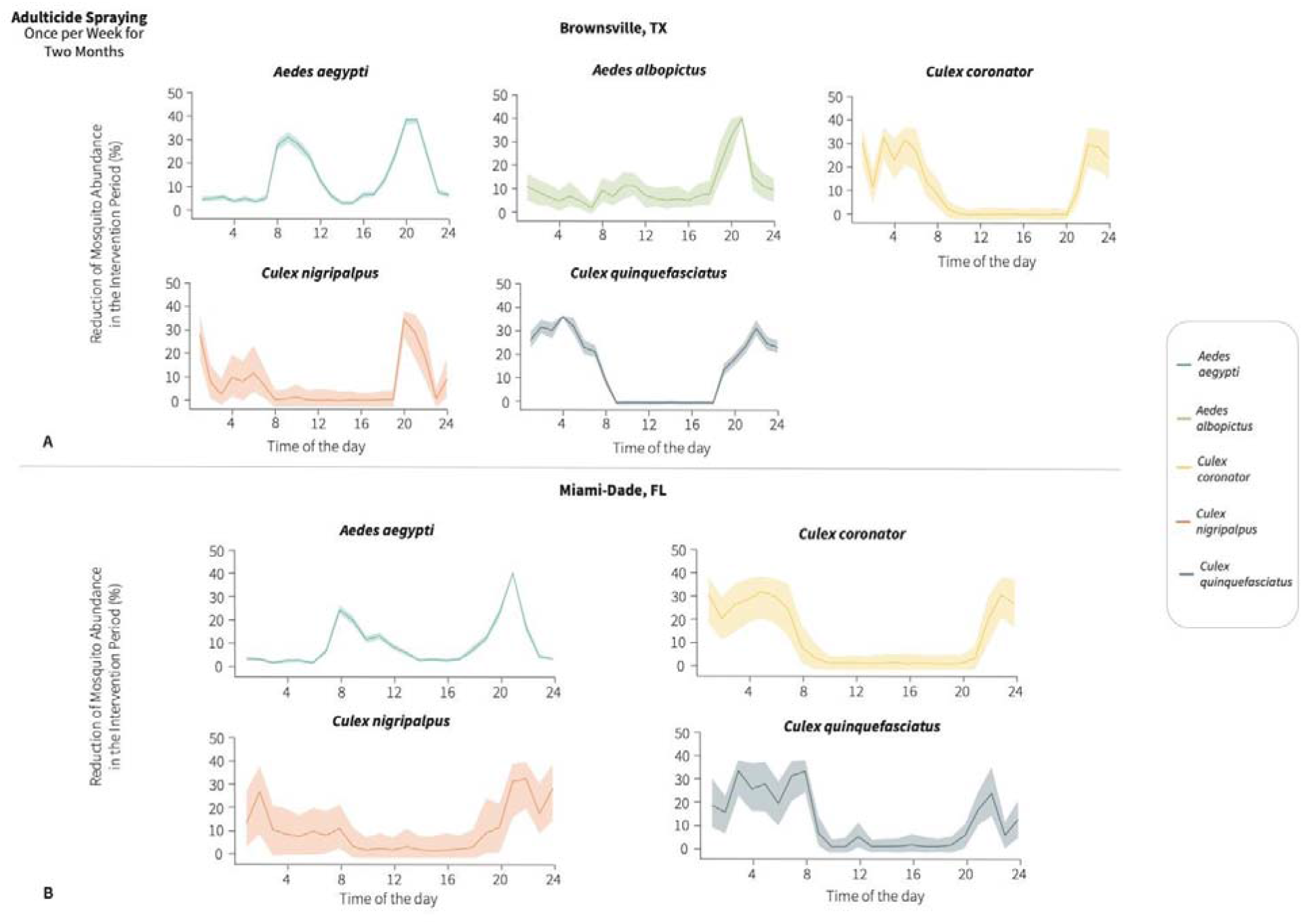
Average reduction in the abundance of populations of vector mosquito species during adulticide interventions comprised of weekly adulticide spraying for two months at different hours of the day in (A) Brownsville, Texas; and (B) Miami-Dade, Florida. Solid line = average number of mosquitoes; Light color highlights = 95% Confidence Interval.

